# Insect pollination is an ecological process involved in the assembly of the seed microbiota

**DOI:** 10.1101/626895

**Authors:** Alberto Prado, Brice Marolleau, Bernard E. Vaissière, Matthieu Barret, Gloria Torres-Cortes

**Affiliations:** INRA, UR 406 Abeilles et Environnement, F 84914, Avignon, France; IRHS, Agrocampus-Ouest, INRA, Université d’Angers, SFR4207 QuaSaV, 49071, Beaucouzé, France

**Keywords:** Bee pollination, bacterial transmission, entomovectors, seed-associated microbes

## Abstract

The assembly of the seed microbiota involves some early microbial seed colonizers that are transmitted from the maternal plant through the vascular system, while other microbes enter through the stigma. Thus, the seed microbiota consists of microbes not only recruited from the vascular tissues of the plant, but also from the flower. Flowers are known to be a hub for microbial transmission between plants and insects. This floral-insect exchange opens the possibility for insect-transmitted bacteria to colonize the ovule and subsequently the seed, and to pass then into the next plant generation. In this study, we evaluated the contribution of insect pollination to the seed microbiota through high-throughput sequencing. Oilseed rape (OSR) *Brassica napus* flowers were exposed to visits and pollination by honey bees (*Apis mellifera*) or red mason bees (*Osmia bicornis*), hand pollination, or autonomous self-pollination (ASP). Sequence analyses revealed that honey bee visitation reduced the bacterial richness and diversity, increased the variability in the seed microbial structure, and introduced bee-associated taxa. In contrast, mason bee pollination had minor effects on the seed microbiota. We highlight the need to consider insect pollination as an ecological process involved in the transmission of bacteria from flower to seeds.

**IMPORTANCE:** Insect pollinators and flowering plants have a very old mutualistic relationship in which animal mobility is used for the dispersal of pollen. The pollination services provided by insects are extremely important to many natural plant populations as well as agricultural crops. Here we show that while visiting flowers, insect pollinators can disperse bacteria that are able to colonize the developing seed via the flower. Hence, insect pollination participates in the assembly of the seed microbiota, the inoculum for the next plant generation. This novel insight has important implications in terms of re-assessing pollinator services by including microbe transfer.

## INTRODUCTION

In nature, plants live in close association with a wide diversity of micro-and macro-organisms, both inside and outside their tissues. Microbes play essential roles in plant growth and development, affecting plant biomass or disease resistance (Berendsen et al. 2012; Santhanam et al. 2015; Sugiyama et al. 2013). Although numerous studies have been focusing on the analysis of microbial assemblages associated to different plant organs (Bulgarelli et al. 2013, Junker et al. 2011), little is known about tripartite interactions between plants, their microbiomes and other multicellular organisms, like pollinators. Insect visitors acquire and deposit microorganisms onto flower surfaces during nectar and pollen collection (Adler et al. 2018; Aizenberg-Gershtein et al. 2013; de Vega et al. 2013; Ushio et al. 2015), thus shaping the flower microbiota (Aleklett et al. 2014; Manirajan et al. 2016; McFrederick et al. 2017). Since the flower microbiota serves as one of several inocula for the plant ovule, and hence for the seed (Truyens et al. 2015), it is possible that by affecting the microbial community of the flower (including pollen), pollinators could modify the seed microbiota.

The role of insect vectors in the dispersal of bacteria and fungi to roots, stems, leaves, flowers, and fruits is well documented (Perilla-Henao et al. 2016; Shikano et al. 2017), while their role in the microbial assembly of the seed has yet to be described. During the seed-to-seed development cycle, some early microbial seed colonizers are transmitted from the mother plant to the ovule through the vascular system (internal transmission) while others colonize the ovule through the stigma (floral transmission). Other microbes are subsequently incorporated to the seed via external transmission, due to the contact with microorganisms present on fruits, flowers or threshing residues (Maude, 1996). Thus, the assembly of the seed microbiota is a complex process, including microbes recruited not only from the vascular tissue of the plant, but also from the floral microbiota. By affecting floral traits, microorganisms inhabiting the flower can have beneficial or detrimental consequences for the reproductive success of the plant (Aleklett et al. 2014; Canto et al. 2008; Herrera et al. 2013). However, it is unknown whether microbes inhabiting the flower can be incorporated via the floral pathway and affect seed traits. It is therefore important to understand the drivers in the assembly of seed microbiota.

Previous community-profiling approaches performed on the seed microbiota of various plant species have identified a range of bacterial taxa that could be potentially insect-transmitted (Adam et al., 2018; Bergna et al., 2019; Barret et al. 2015, Klaedtke et al., 2016; Leff et al., 2017, Links et al., 2014, Rezki et al., 2016; Rybakova et al., 2017, Rodrigues et al., 2018; Truyens et al. 2015). For instance, bacterial taxa affiliated to the *Enterobacteriaceae*, such as the ubiquitous *Pantoea agglomerans*, are found both in flowers, seeds and insect visitors such as the honey bee *Apis mellifera* (Loncaric et al. 2009; Rezki et al. 2018). In addition, another *Enterobacteriaceae* species, *Rosenbergiella nectarea*, isolated from the nectar of different plant species (Halpern et al. 2013), has also been detected in seeds (Torres-Cortes et al. 2018), indicating a possible bacteria transmission to the seed through the floral pathway by insect pollinators. However, it is currently unknown if insect pollinators participate in shaping the structure of the seed microbiota.

In this study, we apply metabarcoding approaches to uncover the contribution of insect pollinators to the seed microbiota and to the transmission of seed-associated microorganisms. Since bees (*Apoidea*: *Anthophila*) are one of the most important pollinators and harbor bacteria that are shared with flowers (Mcfrederick et al. 2017), we have examined the effect of bee pollination on the seed microbiota of oilseed rape (OSR; *Brassica napus*) by performing pollination exclusion experiments. Our results show that bee pollination participates in the microbial assembly of the seed by reducing the bacterial richness and diversity, increasing the variability amongst plants (beta dispersion) and introducing bee-associated taxa. Collectively, these data suggest that insect pollination is an ecological process involved in the assembly of the seed microbiota.

## RESULTS

The effect of bee pollination on OSR seed-associated microbial assemblages was assessed during two years (2017 and 2018) on two different plant lines: i) a male fertile F1 hybrid (MF; cv ‘Exocet’) that was bagged to exclude insects and be autonomously self-pollinated or left open to bee visits with two pollinator species, the domestic honey bee (*Apis mellifera)* and the red mason bee (*Osmia bicornis)*; and ii) its male sterile parent, thereafter referred to as MS line, that does not produce pollen and was either hand-pollinated with pollen from many different plants of the MF line or exposed to honey bee visits and pollination. In total, 198 samples corresponding to bees, nectar, pollen and seeds were analyzed by 16S rRNA gene amplicon Illumina sequencing **(Supp. Table 1**).

**Table 1.** Results of the constrained analysis of principal coordinates. Proportion of variance explained by the indicated variable on the different data sets. The proportion of constrained inertia, F-values and *p*-values were calculated through a canonical analysis of principal coordinates followed by PERMANOVA. Seeds issued from autonomous self-pollination and bee pollination are referred to as ASP and BP seeds, respectively.

| Data set name | Year | Data set material | Explanatory variable | Samples | Proportion of constrained inertia (%) | F-value | *p*-value |
| --- | --- | --- | --- | --- | --- | --- | --- |
| All samples | 2017<br>2018 | ASP and BP seeds, nectar, pollen, bees | Year | 198 | 5.7 | 11.45 | 0.0001 |
| Experiment 2017 | 2017 | ASP and BP seeds, nectar, pollen | Material | 16 | 35.54 | 3.584 | 0.0002 |
| Experiment 2017 male fertile line | 2017 | ASP and BP seeds | Pollination mode | 10 | NA | 1.19 | NS |
| Experiment 2017 male sterile line | 2017 | ASP and BP seeds | Pollination mode | 12 | NA | 1.073 | NS |
| Experiment 2018 | 2018 | ASP and BP seeds (honey bee and mason bee), nectar, pollen | Tunnel | 134 | 4.5 | 6.22 | 0.0001 |
| Honey bee experiment 2018 | 2018 | ASP and BP seeds, nectar, pollen | Material | 69 | 9.3 | 3.28 | 0.0001 |
| Honey bee experiment 2018 male fertile line | 2018 | ASP and BP seeds | Plant ID | 18 | NA | 0.91 | NS |
| Honey bee experiment 2018 male fertile line | 2018 | ASP and BP seeds | Pollination mode | 18 | 12.3 | 2.23 | 0.001 |
| Honey bee experiment 2018 male sterile line | 2018 | ASP and BP seeds | Pollination mode | 21 | NA | 1.29 | NS |
| Mason bee experiment 2018 male fertile line | 2018 | ASP and BP seeds | Plant ID | 20 | NA | 1.33 | NS |
| Mason bee experiment 2018 male fertile line | 2018 | ASP and BP seeds | Pollination mode | 20 | NA | 1.14 | NS |

### Experimental design and sequence analysis

To investigate if bee pollination services has an additional effect on the microbial assemblage of OSR seeds, bee and plant samples were collected and submitted to bacterial community profiling analyses. Pollinator exclusion experiments were carried out inside two insect proof tunnels. Prior to the experiment, all inflorescences were covered under gas permeable osmoflux bags to exclude any contact with insects. During the experiment, half of the inflorescences on each plant were uncovered and exposed to bee pollination (seeds issued from this treatment will be referred to as BP seeds). The other half of the inflorescences were left covered. These un-touched flowers produced seeds resulting from autonomous self-pollination, hereafter referred to as ASP seeds. Inflorescences of the MS line were pollinated by hand using a paint-brush with pollen of many different plants (HP seeds). In this way, ASP, BP and HP seeds, as well as nectar, pollen and bees were collected and their DNA was extracted. Amplicon libraries of the V4 region of 16S rRNA were constructed afterwards and sequenced to characterize the bacterial assemblages present in the samples. The raw sequencing data was first resolved into Amplicon Sequence Variants (ASVs) using the software DADA2 (Callahan et al. 2016). After quality filtering, chloroplast removing and taxonomic affiliation, a total of 6.3 million reads were classified in 2,764 ASVs. According to rarefaction curves, most samples reached an asymptote at 12,000 sequences and were rarefied at this value (**Supp. Figure S2**). This analysis indicated that we were able to characterize most of the diversity found in the seed samples with an average of 95.7 ASVs per seed sample in the MF line and 67.4 ASVs per sample in the MS line.

### Taxonomic composition of bee, pollen, nectar and seed microbiota

Analysis of the ASV taxonomic affiliation showed that *Proteobacteria* and *Firmicutes* were the dominant *phyla* in honey bee samples (**Supp. Figure S3**). These phyla contain the main taxa of the “core gut” microbiome in honey bee workers such as *Frischella*, *Gillamella*, *Snodgrassella*, *Lactobacillus* and *Buchnera* spp. (Engel et al. 2012). Concerning OSR pollen, samples were also dominated by *Firmicutes* and *Proteobacteria* in both years. However, we observed very contrasting results for nectar samples between years (**Supp. Figure S3**). Nectar samples from 2017 were more diverse than the ones from 2018, which were dominated by the genus *Acinetobacter*. This taxa was also found in very high abundance in the seeds (see below).

In the case of microbial assemblages associated to seeds, the most common bacteria belonged to the *Actinobacteria*, *Bacteroidetes*, *Firmicutes* and *Proteobacteria* (Figure 1a, 1c), which is in agreement with other microbiome studies performed on OSR seeds (Rybakova et al. 2017). Seed samples from 2018 were dominated by the genera *Acinetobacter* and *Pantoea* (around 80% of the reads were affiliated to these two genera; Figure 1c). Honey bee pollination changed significantly (logarithmic LDA score higher than 2.5) the abundances of 3 ASVs in the seeds, while mason bee pollination changed the abundances of 15 ASVs (Figure 1d; **Supp. Table 3 and 4**).

**Figure 1.**
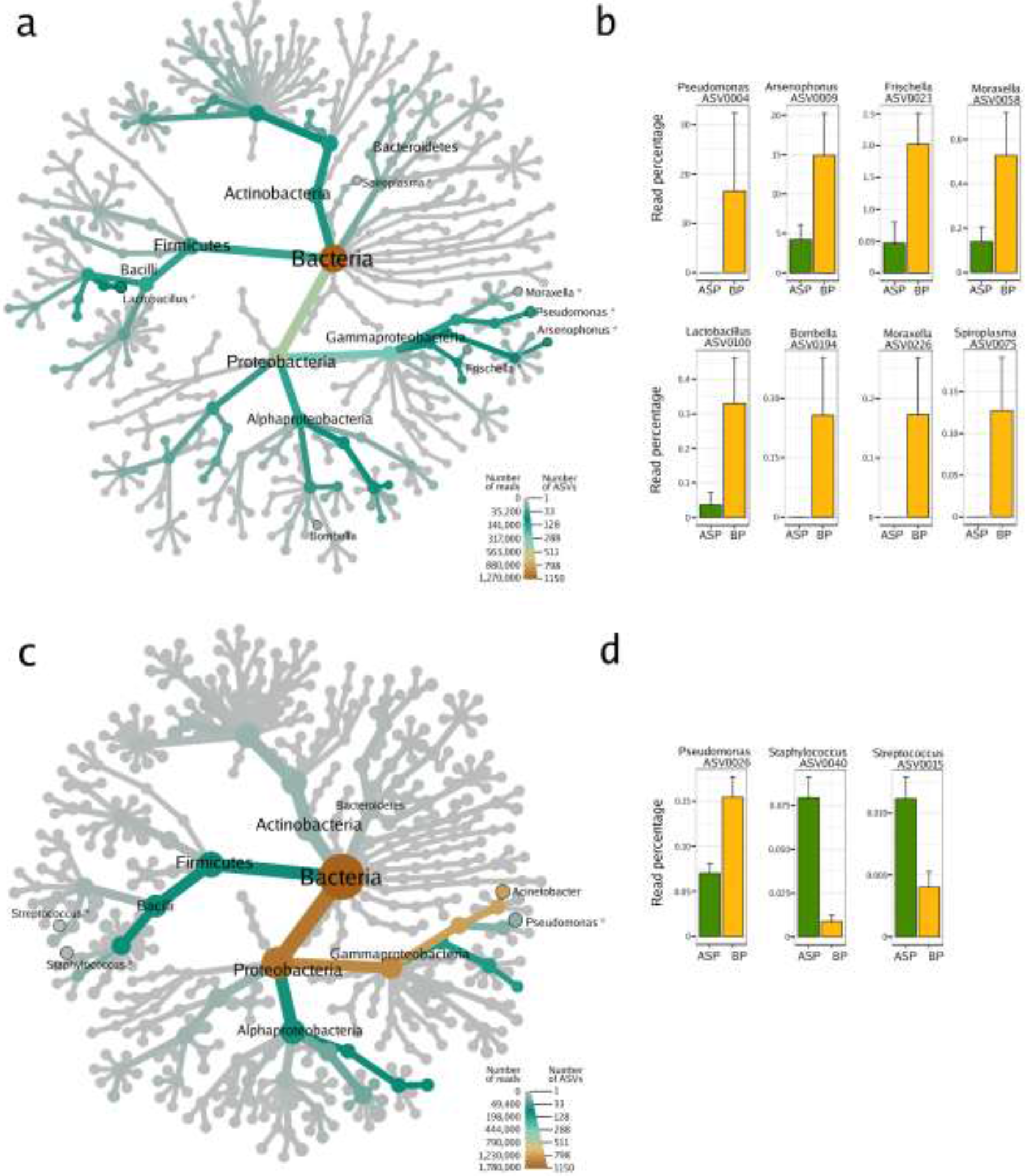
Microbial composition of oilseed rape seed samples issued from honey bee pollination or autonomous self-pollination. Heat trees show the microbial composition of the seeds samples harvested in 2017 (**a**) and 2018 (**c**). The size of the nodes refers to the number of ASVs of known identity and the color of the nodes and edges represent the ASV read abundance. Asterisks (*) indicate the taxonomic affiliation of ASVs with significant changes in relative abundance (according to Linear Discriminant Analysis Effect Size; LefSe) in relation to the pollination mode. ASVs with significant changes in relative abundance are displayed on the right part of the figure (**b**, **d**). BP: honey bee-pollination; ASP: autonomous self-pollination.

In the seed samples collected during 2017, *Acinetobacter* and *Pantoea* were not the most abundant taxa; the genera *Sphingobium*, *Pseudomonas*, *Lactobacillus* and *Gillamella* were the most predominant instead. Interestingly, honey bee pollination affected the relative abundances of 7 ASVs (Figure 1b; **Supp. Table 2**), 4 of which are specifically associated with honey bees and a part of the bee core microbiota (Figure 1a). *Arsenophonus* (Yañez et al. 2016), *Frischella* (Engel et al. 2013), *Spiroplasma* and *Lactobacillus* are bee-associated taxa and were significantly more abundant in seeds issued from BP as opposed to those issued from ASP. Specifically, the ASV0075 showed 99% identity with the honey bee pathogen *Spiroplasma apis* strain B31^T^ (Ku et al. 2014) and the ASV0100 showed 100% identity with *Lactobacillus mellis* strain H1HS38N, a symbiotic bacterium inhabiting the bee stomach (Olofsson et al. 2014). The other two remaining taxa enriched in bee-pollinated samples belong to the genus *Moraxella*, which is highly abundant in nectar. The acetic acid bacteria of the genus *Bombella*, which are found in the gut of bumble bees and honey bees (Li et al. 2015), were only present in the bees themselves and in the seeds issued from bee pollination. However, due to its low relative abundance, the *Bombella* ASV did not achieve the 2.5 fold change criteria. These results suggest that insect pollinators can transfer bacteria (*Arsenophonus*, *Frischella*, *Spiroplasma*, *Lactobacillus* and *Bombella*) from the insect itself or from the nectar collected to the seed through the floral pathway.

### Seed microbial alpha diversity is modified by honey bee pollination

To further investigate how bee pollination affects the structure of seed microbial assemblages, bacterial richness and diversity indexes were assessed for the seed-associated assemblages in all the different treatments (Figure 2). To compare the alpha diversity in seed samples, non-parametric Wilcoxon rank-signed tests were performed on the rarefied data set. No statistically differences were found in alpha diversity in the 2017 seed samples (N=11). In contrast, variations of ASV richness and diversity were observed between seed samples in relation to the pollination mode in 2018 (N=53, Figure 2a). Bacterial richness significantly decreased in seeds issued from honey bee pollination (p<0.0001; Figure 2a). However, we do not observe the same trend in the seeds issued from mason bee pollination (Figure 2a). These discrepancies observed in the two types of pollinators used in this study are probably due to differences in the foraging intensity of these insects. Indeed, honey bees visited the flowers more intensively (~ 1 visit per flower every 2 min) than mason bees (~ 1 visit per flower every 10 min). Moreover, while the honey bee colony used in these experiments consisted of a population of ca. 5000 worker bees with a population of foragers of at least 1000 individuals, only 200 female and male cocoons of mason bees were introduced. It is also noteworthy that the visits to the experimental flowers during the mason bee experiment was done mostly by males that were searching for nectar between matings.

**Figure 2.**
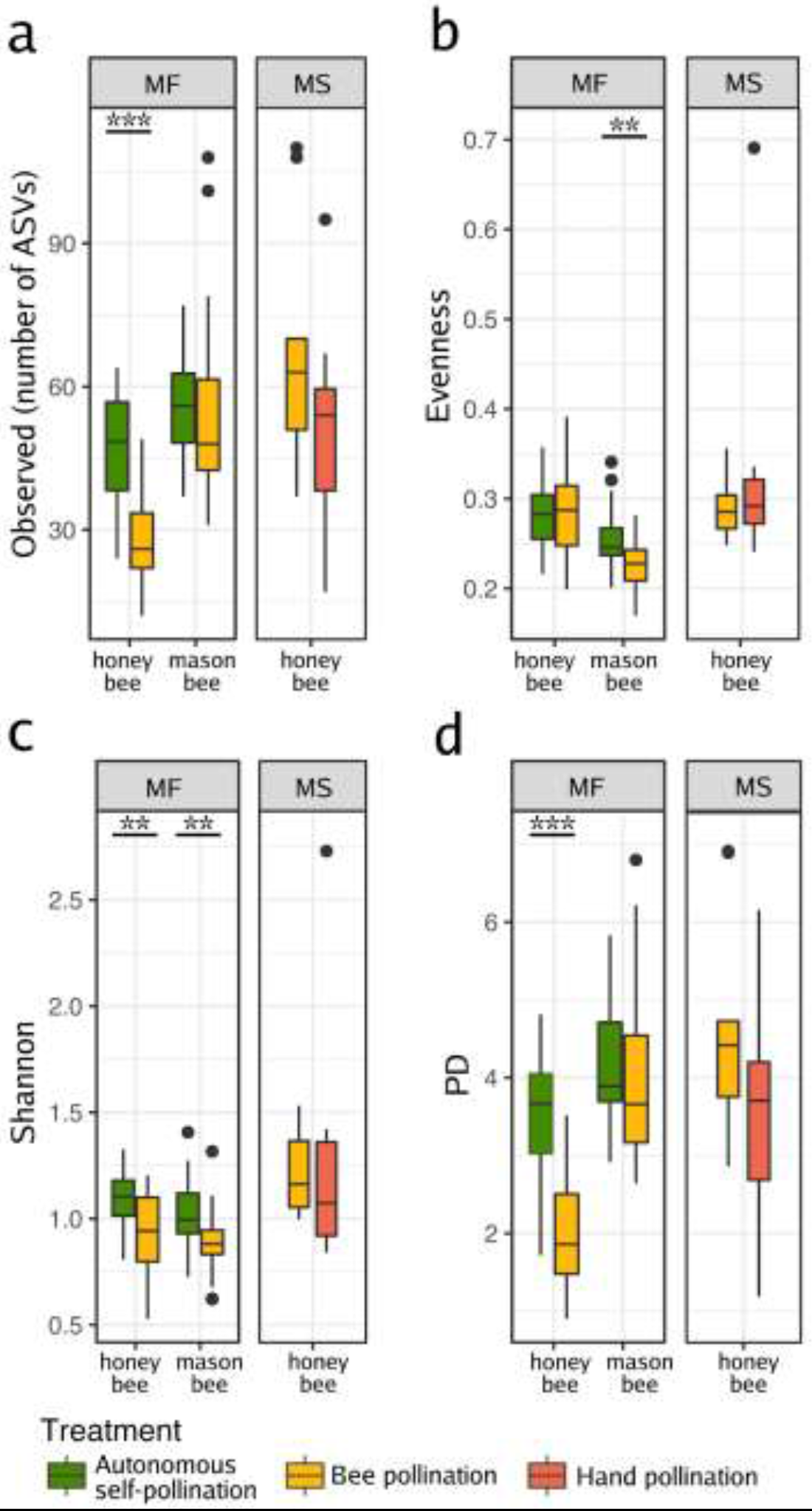
Changes in microbial richness and alpha diversity between seed samples. Observed richness (**a**), evenness (**b**) and diversity (Shannon and Faith’s PD phylogenetic diversity; **c**, **d**). The indexes were estimated in seed samples harvested from oilseed rape male fertile plants pollinated by bees or left for autonomous self-pollination. Additional indexes were calculated on seed samples harvested from a male sterile line that was hand- or insect-pollinated. Wilcoxon rank-signed tests were performed to assess the effect of pollination on richness and alpha diversity. Asterisks denote statistically significant differences between conditions considered as a *p-value*<0.05 (*), a *p-value*<0.01 (**), and a *p-value*<0.001 (***). Richness, evenness and Shannon diversity were assessed with the number of ASVs rarefied at 12,000 sequences per sample. MF: male fertile line; MS: male sterile line.

Seeds issued from hand pollination on the MS line (HP seeds), did not differ in bacterial richness from those issued from bee pollination, suggesting that the decrease in richness (Figure 2a) could be partly explained by the amount of pollen deposited on the stigma and not by the selection of insect-associated taxa. When checking the evenness in the samples, we observed a significant reduction due to mason bee pollination. Indeed, *Acinetobacter* is the dominant taxa in seeds issued from mason bee pollination.

To assess the impact of bee pollination on the diversity of the seed microbiota, different diversity indexes were calculated. As observed with richness, there was a significant reduction in bacterial diversity due to honey bee pollination (Shannon and Faith’s PD phylogenetic diversity; p=0.003 and p<0.0001 respectively; Figure 2c-d). In this case, we observed a similar trend in the diversity of the microbial assemblages associated to the seeds issued from mason bee pollination (Figure 2c), where the diversity calculated with the Shannon index was also reduced (p=0.001). Altogether, these results suggest that bee pollination decreases the diversity of the microbial assemblages associated with seeds.

### Effect of bee pollination on the structure of the seed microbiota

Similarity in community composition between samples was then estimated through PCoA ordination of unweighted UniFrac distances (Table 1 and Figure 3 for 2018 data). The relative contribution of pollination was then investigated through canonical analysis of principal coordinates (CAP) followed by PERMANOVA (Table 1). The bacterial composition differed between plant materials (seeds, nectar and pollen; Figure 3a), nectar and seeds clustering together in the ordination plot and separated from pollen. The type of plant material explained 35.54 and 9% of the variation in microbial composition in 2017 and 2018 respectively (Table 1; *p*=0.0002 and *p*=0.0001). As expected, microbial assemblages associated to bees were distinct from the ones associated to plant tissues. Moreover, microbes associated with the surface of the highly social honey bees were very different to the microbes living inside the bee gut (Engel et al. 2012). However, this is not the case for the solitary mason bee, which harbored similar microbial assemblages on its surface and gut (Figure 3b).

**Figure 3.**
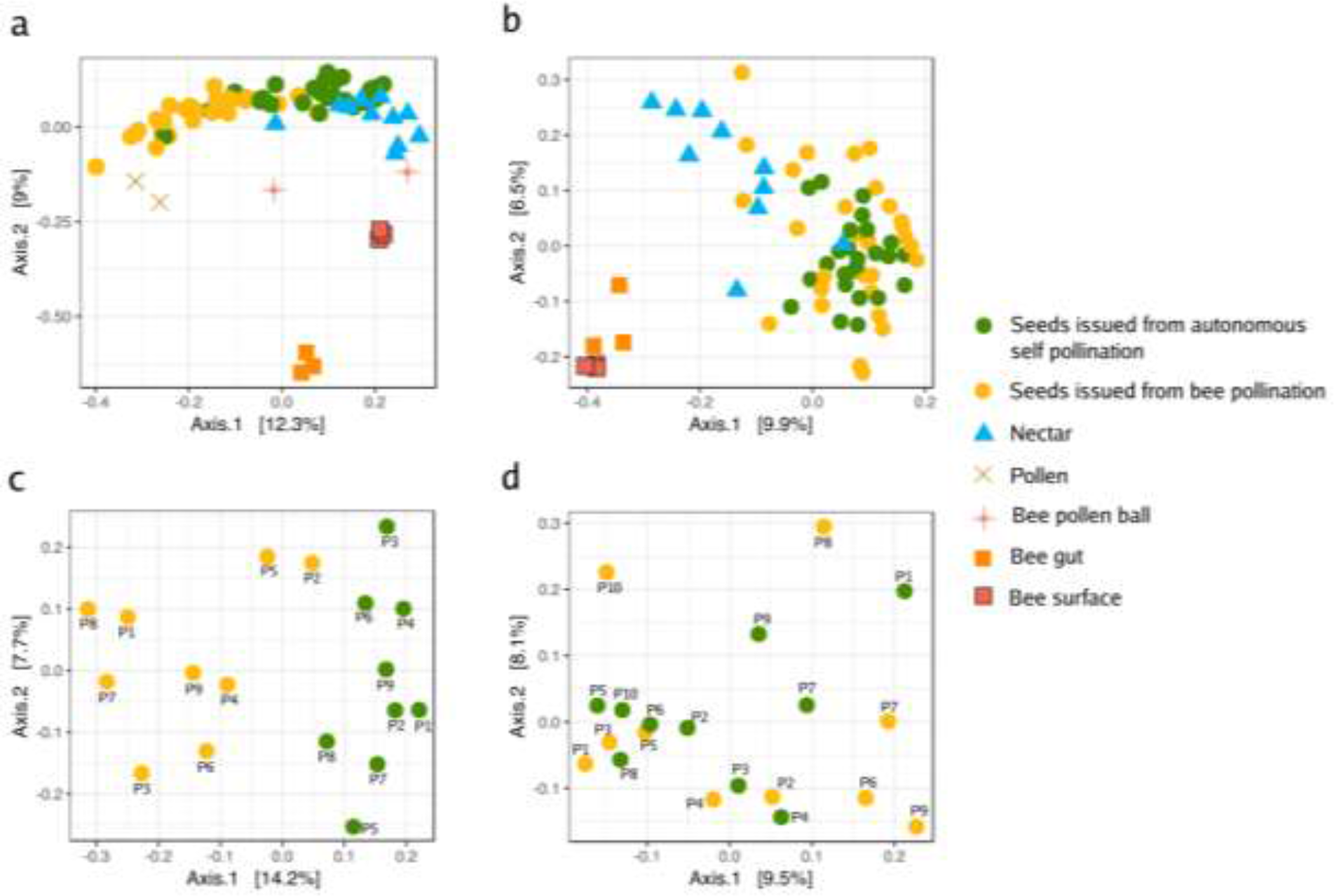
Ordination of unweighted UniFrac matrices with principal coordinate analysis (PCoA) showing variation in microbial composition. PCoA plots show the ordination of all samples from the honey bee (**a**), and the mason bee (**b**) experiment. Seed microbial assemblages of each plant (pooled samples) are represented in panel (**c**) for HB pollination, and in panel (**d**) for MB.

Differences in microbial composition of ASP and BP seeds were not statistically different in 2017 (Table 1). However, in 2018, flowers pollinated by honey bees produced seeds with significant changes in microbial composition compared with those from ASP seeds, showing that honey bee pollination can change the composition of the seed microbiota (Figure 3c). Moreover, these BP seed microbial assemblages diverge from those of ASP seeds and are closer to those of pollen (Figure 3a). According to CAP analyses, honey bee pollination explained 12.3% (*p* = 0.001) of the variation in bacterial community composition in the seed samples in 2018. In the experiments performed in 2017, we observed a similar trend, although the change was not statistically different due to the low size of the sample (Table 1). In contrast, we did not observe changes in the microbial composition of the seeds issued from flowers pollinated by mason bees, probably due to the reduced visitation rates observed during the experiment.

During both years, the bacterial composition of the seeds produced by honey bee pollination did not differ from that of HP seeds on the MS line (Table 1). Ordination plots did not show a segregation between the seeds from the honey bee and the hand pollination treatments. In agreement with these results, the constrained analysis of principal coordinates revealed that the pollination mode of the sterile line had no significant effect on seed-associated microbial assemblages. This fact suggests that the differences observed between seeds that resulted from honey bee pollinated flowers and seeds that came from ASP flowers could be due to the amount and the varied origin of the pollen, and not necessarily to the insect associated taxa.

### Pollination by honey bees increases microbial beta dispersion among seeds

The PCoA (Figure 3c) showed that following honey bee pollination, the bacterial assemblages associated to BP seeds were more variable than those of the ASP seeds. Our experimental design involved having two pollination modes represented within each plant, which allowed for the comparison of the variability in the bacterial composition of seeds issued from those pollination modes. We compared the distances of each plant in the principal coordinate space (created using the unweighted UniFrac distances) to the group (pollination mode) centroid (Anderson et al., 2006; Vannette et al. 2017). In this way, we assessed the beta dispersion as the variability of the ASV composition at the plant level (inter-plant variation, n=9, n=10 & n=10 for honey bee, mason bee and HP on the MS line respectively). We did not observe statistically significant changes in distance to the centroid (beta dispersion) between ASP and BP seed microbial assemblages in the mason bee treatment. No differences in beta dispersion were detected between honey bee and hand pollinated seed microbial assemblages in the MS line. However, we did observe that the beta dispersion was significantly higher in seed communities that were exposed to honey bee pollination in contrast to those that came from flowers with ASP (Wilcoxon rank-signed test, *p* = 0.02; Figure 4).

**Figure 4.**
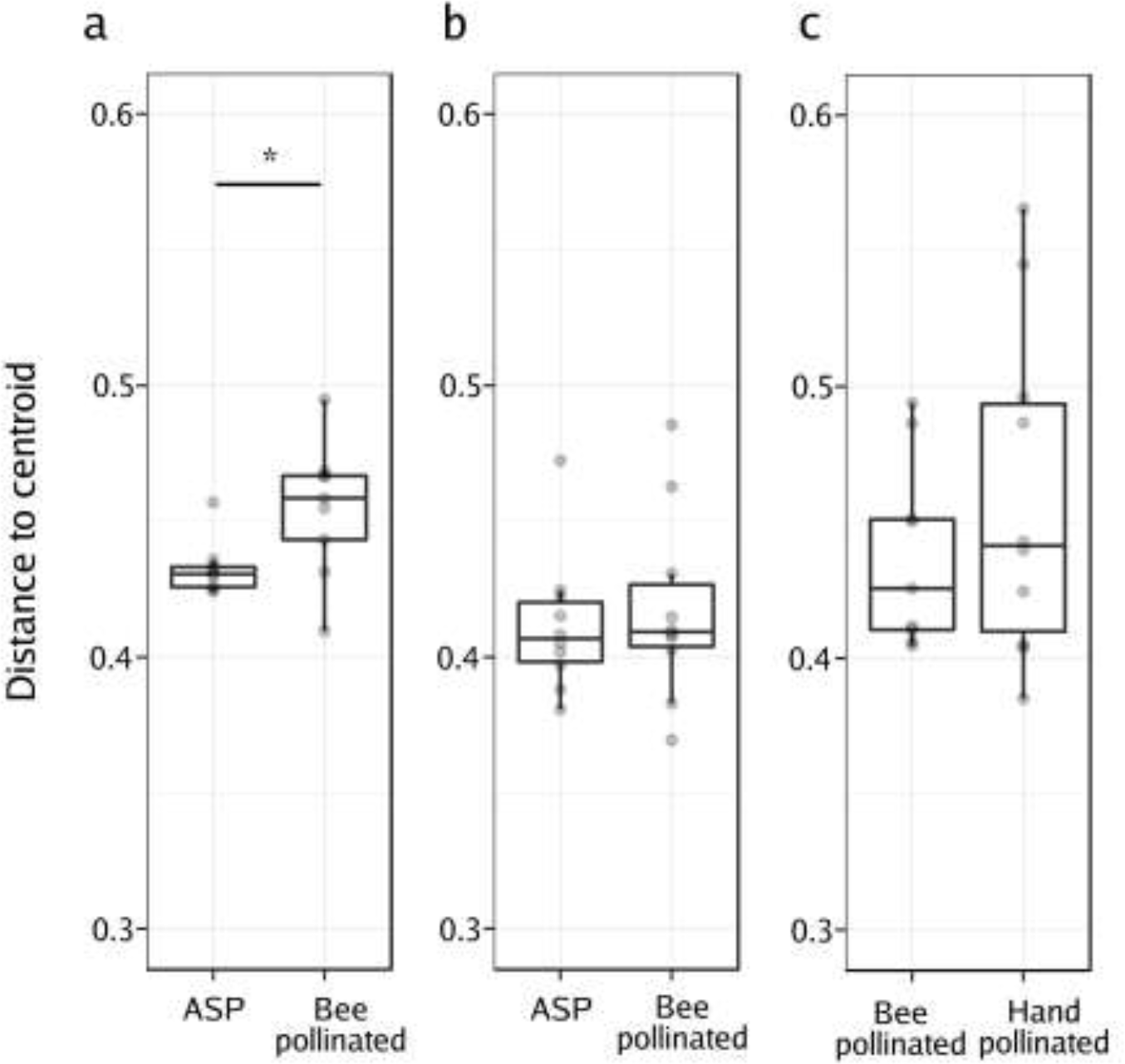
Effect of the pollination mode on the structure of seed microbial assemblages. Analysis of the multivariate homogeneity of group dispersions (variances). Boxplots represent the distance to the centroid of seed-associated microbial communities of male fertile (a, b) and male sterile plants (c) submitted to different modes of pollination. **a)** Distance to the centroid of seeds issued from male fertile plants subjected to autonomous self-pollination (ASP) or honey bee pollination, **b)** distance to the centroid of seeds issued from male fertile plants subjected to ASP or mason bee pollination, **c)** distance to the centroid of seeds issued from male sterile plants subjected to honey bee pollination or hand-pollinated with pollen of different plants. Wilcoxon signed-rank tests were performed to assess the effect of the pollination mode on the distance to the centroid. Asterisks denote statistically significant differences between conditions considered as a *p-value*<0.05 (*).

## DISCUSSION

Currently, little is known about the neutral or niche-based ecological processes involved in the assembly of the seed microbiota. Niche processes, like selection by the environment, have been shown to shape the structure of seed-associated fungal assemblages (Klaedtke et al. 2015). Furthermore, host-filtering processes have been shown to be involved in the assembly of oilseed rape (Rybakova et al. 2017) and tomato (*Solanum lycopersicum*) seed microbiomes (Bergna et al. 2018). On the other hand, neutral processes related to ecological drift have been described as important drivers of the structure of the bacterial communities of radish (*Raphanus sativus*) seeds (Rezki et al. 2018). Taken together, these pioneer studies suggest that seed-associated microbial communities consist of a few dominant taxa that are probably niche-selected, and multiple scarce taxa whose distributions could be explained by neutral processes. The present study supports the hypothesis that insect pollination is a neutral ecological process (microbial dispersal) that participates in the dispersal of microbes between flowers and seeds and influences the assembly of the seed microbiota. Our results show that *Apis* pollination can change diversity of seed-associated bacterial assemblages and in some cases introduce bee-associated taxa.

The proximity between a floral structure and the developing seed suggests that microbes associated with the flower could eventually colonize the developing seeds through the floral pathway. The floral pathway for the colonization of the seed tissue is less specialized than the vascular (internal) pathway. Indeed, plant-pathogenic bacteria can colonize both host and non-host seeds via the floral pathway (Darsonval et al., 2008). Thus, the mechanisms dispersing microbes into flowers (like insect pollination) could have important implications for the transmission of non-specialized microbes to the developing seeds. Previous studies have already shown that insect pollination modifies the composition of microbial assemblages associated to flowers (Ushio et al 2015), nectar (Aizenberg-Gershtein et al. 2013; de Vega et al. 2013; Vannette et al. 2017) and pollen (Manirajan et al. 2016); however, virtually nothing is known about the effect of insects pollinators on plant seed microbial communities. According to our data, the changes observed in the seed microbiota were mostly associated with the honey bee (*A. mellifera*) and very mild effects were observed with the red mason bee (*O. bicornis*). During our experiments, the bee density between the two bee species differed considerably (honey bees being an order of magnitude more abundant) and we observed a strong difference in visitation rates. These differences are reflected in our results on the seed microbiota, suggesting that the amount of contact between the insect and the flower might be a determining factor in the effect of the insect on the seed microbiota. Indeed, the time that a bee spends foraging on a flower is correlated with the transmission of bee pathogens (Adler et al. 2018). The potential role that foraging behaviour (especially the foraging intensity and the handling time, i.e. time spent per flower) has on the seed microbiota should be assessed in order to shed light on the role that different pollinators might have in the transmission of seed-borne diseases (Aleklett et al. 2014; Truyens et al. 2015).

The microbial assemblages associated with seeds examined in this study are dominated by members of the *Actinobacteria*, *Bacteroidetes*, *Firmicutes* and *Proteobacteria* phyla (Figure 1). These phyla were also prevalent in other studies of the oilseed rape seeds (Rybakova et al. 2017). The taxonomic profiles were very different between the two years. This is not so surprising since biotic factors (such as the type of soil, the climatic conditions, and also the field management practices) have a strong influence on the seed microbiota (Klaedtke et al. 2016). *Acinetobacter* was the dominant genus of the seeds and the nectar in 2018. This is a widespread genus in animals, plants and the environment (Doughari et al. 2011). Specifically in plants, *Acinetobacter* spp. have been described as plant-growth-promoting bacteria (Suzuki et al. 2014). In our 2018 experiments, this taxon was already present in samples of pollen and nectar prior to the pollination exclusion experiments. It is thus possible that *Acinetobacter* acted as a pioneer taxon in seeds exerting a strong priority effect in the final microbial composition of the seed.

While mason bee carries pollen in its abdominal scopa, the honey bee aggregates pollen grains by adding regurgitated nectar or diluted honey to transport it to its nest in the corbiculae of its hind legs. The addition of nectar rises the humidity of the pollen grain, causing it to swell and expose the intine (Human & Nicolson 2008). The taxonomic profiles of the 2017 seed samples showed that honey bee pollination changed the seed microbiota by increasing the abundance of bee- or nectar-associated taxa. It is then tempting to speculate that the way the honey bee processes pollen allows for the transmission of bee-associated bacteria to the seed. Indeed, *Spiroplasma*, *Lactobacillus*, *Arsenophonus*, *Frischella* and *Bombella* are insect symbiotic bacteria living in the bee gut (Corby-Harris et al. 2014; Engel et al. 2013; Yanez et al. 2016; Yun et al. 2017) and were more abundant (and in some cases, only present) in seeds issued from BP as opposed to ASP. These results illustrate the possibility of insect-transmitted bacteria colonizing the seed. Interestingly, two of the identified ASVs affiliated to *Spiroplasma* and *Arsenophonus* could act as bee pathogens, suggesting that plants could act as a reservoir for bee pathogens.

In our experimental design, the effect of bee pollination on the seed microbiota was assessed in two different plant genotypes: a male fertile line and a male sterile line that does not produce pollen. Our results on the male sterile line show that hand pollination did not differ from bee pollination. This would suggest that the diverse origin (many source plants) and copious amount of pollen delivered either by a bee or by a paintbrush have similar effects on the seed microbiota. On the contrary, the reduced amount and single origin of the pollen of ASP causes significant differences to the seed microbiota as compared with that of the seeds resulting from bee pollination. It is known that the amount of pollen delivered to the stigma has an impact on the germination rates of the pollen grains: small pollen populations germinate poorly. This population effect is partly explained by the availability of certain growth factors such as calcium ions (Brewbaker & Kwack 1963), flavonols (Taylor & Hepler 1997) and phytosulfokine-alpha (Chen et al. 2000). The greater availability of nutrients presented by large pollen populations might also provide nutrients for bacteria. The availability of nutrients fosters competition between the microbial taxa, which could explain the observed decrease in species richness observed in honey-bee-pollinated seeds.

Alternatively, the decrease in alpha diversity might be related to the period during which the stigma is receptive. Bacteria are generally unable to actively penetrate the plant tissue and rely on openings such as wounds, stomata, lenticels and nectarhodes (Gimenez-Ibanez et al. 2010). A possible route to enter the seed is provided by the stigma surface through which bacteria can penetrate (Compant et al. 2011; Truyens et al. 2015), possibly during the penetration of the pollen tubes. Following flower opening, oilseed rape flowers require on average ~13h of exposure to pollinators to complete their sexual function (Bell & Cresswell 1998). The removal of pollen triggers flower senescence (Bell & Cresswell 1998), so in our experiments the flowers visited by bees were likely to have senesced faster than the flowers left to self-pollinate autonomously. The increased duration of flower longevity in the ASP treatment may have facilitated the entry of bacteria, explaining then the higher diversity of the seed microbial communities. Future experiments should aim at disentangling how insect pollination and flower longevity affect the seed microbiome.

Honey bee pollination enhanced the variation in the structure of seed bacterial assemblages (Figure 4a). This result is in agreement with Vanette *et al.* (2017) who reported that pollinators increase dispersal of microorganisms and ultimately enhance dissimilarity between nectar microbial assemblages. These findings could be explained by a high stochasticity in the order in which bacterial species arrive (stochasticity of microbial dispersal), and due to strong priority effects, the composition of the assemblages can diverge (Vannette et al. 2017). Pollination by *Apis* would then favor the arrival of new species to the flower and seed, increasing the variability of the seed-associated microbial communities. Our data could serve as a foundation for additional experiments that target directly the effect of priority effects on beta dispersion and on the assembly of the seed microbial communities.

## Conclusions

This study aimed to uncover the contribution of bee pollination to the seed microbiota. We have identified differences in richness, diversity and species composition in seeds issued from bee pollination, as compared to those from flowers with autonomous self-pollination. Our findings with two different bee species suggest that foraging behaviour (foraging rates/intensity) might mediate the insect’s effect on the seed microbiota. Additionally, the amount and origin of the pollen might also have an effect. These results provide novel insights about determinants involved in the transmission of bacteria from flower to seeds, and have important implications in terms of re-evaluating the services provided by pollinators, which could include microbe transfer.

## MATERIALS AND METHODS

### Pollinator exclusion experiments

Pollinator exclusion experiments were performed inside two 22 × 8 m insect proof tunnels at the INRA research station at Avignon during 2017 and 2018. Inside the tunnels, seeds of oilseed rape (*Brassica napus*) were sown in the soil in four 18 m long rows (20 plants per row). Two winter oilseed rape lines were sown side by side: a male fertile F_1_ hybrid line ‘Exocet’ and its male sterile parent which does not produce pollen. Winter oilseed rape was chosen because it is a highly self-fertile plant that produces nectar attractive to bees, and because a male sterile line was available. Plants were watered two times a day with an automated water drip system.

In 2017, the pollinator exclusion experiment was only carried out using honey bees on 5 plants of the male fertile line in a single tunnel. In 2018, the pollinator exclusion experiments included 1) honey bees on the male fertile line in one tunnel 2) mason bees on the male fertile line in a second tunnel and 3) honey bees on the male sterile line in the first tunnel. For each experiment, ten plants were chosen as experimental plants based on their homogenous appearance. On each plant, six flowering panicles were marked using flower markers of two different colours (three panicles each) and bagged with hydrophilic plastic bags made of osmolux film (Pantek France, www.pantek-france.fr/agriculture.html). The osmolux bags are gas-permeable, but prevent all contact with insects, even small ones such as thrips (Thysanoptera) (Perrot et al. 2018). On the day of the introduction of the bees into the tunnels, three color-coded panicles were un-bagged and exposed to bee visits (bee pollination treatment), while the three others remained covered (autonomous self-pollination treatment). Foraging bees were allowed to forage freely amongst the experimental plants and those that were not selected for the experiment. On the male sterile line, the panicles that remained bagged during the introduction of the bees were pollinated by hand using a fine paint brush with pollen coming from many flowers (>200) from several plants of the male fertile line (>10). After 48 hours, the bees were removed from the tunnels and the uncovered panicles were re-bagged. All panicles were kept bagged for an additional 48 hours to ensure that no bee was left in the tunnels, at which point all bags were removed.

One tunnel was used to perform experiments using honey bees (*Apis mellifera*) where we introduced a 5-frame hive (adult population ~5000). In the other tunnel, 100 male and 100 female cocoons of the red mason bee (*Osmia bicornis*) were introduced. Male mason bees were highly active visiting flowers and mating with the females during the experiment. The female mason bees were mostly inactive after mating and due to our experimental design, they were removed from the tunnel before they started provisioning their nests.

### Material collection

Prior to the introduction of bees into the tunnels, pollen and nectar samples were collected from bagged flowers and kept at −20 °C until extraction. Pollen was collected by dissecting closed flower buds and separating the anthers. Anthers were dried for 4 h at room temperature in glass Petri dishes. To recover the pollen, the dried anthers were placed in a steel tea ball and vibrated using a Vibri Vario tomato vibrator. Nectar was collected with 2 μl capillary tubes between the base of the anthers and then transferred to 2 ml Eppendorf containers using a pipette bulb. During the experiment, honey bee foragers were captured and their pollen loads removed (in the case of honey bees) and frozen for further analysis. Once mature (2 months after pollination), OSR fruits (siliques) were collected in large paper bags and taken into the lab. In aseptic conditions, seeds were removed from the siliques and placed in small paper bags.

### DNA extraction, amplicon library construction and sequencing

For seed sample preparation, a total of 0.5 g of oilseed rape seeds of each sample were transferred to sterile tubes containing 2 ml of PBS supplemented with 0.05% (vol/vol) of Tween 20. Samples were incubated for 2 h and 30 min at 4°C under constant agitation (150 rpm). In the case of bee samples preparation, to obtain bee surface microbial assemblages, bees were sonicated in 1 ml of PBS buffer with Tween 20 0.05% (vol/vol). After removing the liquid, insect samples were re-suspended in 1 ml of PBS and crushed to recover the microbes living inside the bees. All the suspensions were then centrifuged (12,000 × *g*, 20 min, 4°C) and pellets were stored at −20°C until DNA extraction. Total DNA extraction was performed with the PowerSoil DNA isolation kit (MoBio Laboratories) using the manufacturer’s protocol.

Amplification, purification and pooling for amplicon library construction were conducted following the protocol described in Barret *et al.* 2015. Briefly, for amplicon construction two rounds of PCR were performed. The first round was designed to target the region V4 of the 16S rRNA with the PCR primers 515f/806s (Caporaso et al. 2011). All PCRs were performed with a high-fidelity polymerase (AccuPrime *Taq* DNA polymerase; Invitrogen) using the manufacturer’s protocol and 10 μl of environmental DNA suspension. After amplicon purification, a second round of amplification was performed with 5 μl of purified amplicons and primers containing the Illumina adaptors and indexes. All amplicons were purified, quantified and pooled in equimolar concentrations. Finally, amplicons libraries were mixed with 10% PhiX control according to Illumina’s protocols. Two sequencing runs were performed in this study with MiSeq reagent kit v2 (500 cycles) for the samples of 2017 and MiSeq reagent kit V3 (600 cycles) for the samples of 2018.

### Data Analysis

MiSeq runs were analysed separately. Primers sequences of fastq files were first cut off using Cutadapt 1.8 (Martin 2011). Files were then merged and processed with DADA2 v.1.8.0 (Callahan et al 2016) according to the recommendations of the workflow “DADA2 Pipeline Tutorial”. The workflow was modified in the truncLen parameter to adjust it to the quality of the sequencing run. The 16S rRNA amplicon sequence variants (ASV) resulting from DADA2 were aligned with a naive Bayesian classifier against the Ribosomal Database Project training set 16 database. Statistical analyses were done with Rstudio v3.3 using the R package *phyloseq* v1.24.2 (McMurdie & Holmes 2013). The *Metacoder* R package v 0.3.0.1 (Foster et al. 2017) was used to plot the distribution of ASV, associated with a taxonomic classification in heat trees. Observed taxa richness, evenness, and diversity were calculated on a rarefied dataset at 12,000 reads per sample and differences were assessed by Wilcoxon signed-rank tests. Variances in community composition between the different samples were assessed by unweighted UniFrac distance (Lozupone & Knight 2005). Principal coordinate analysis (PCoA) was used for ordination of UniFrac distances. Permutational multivariate analysis of variance (PERMANOVA, Anderson 2017) was calculated to investigate the effect of pollinators on microbial community profiles as implemented by the package *vegan* v2.5-3 in R. Moreover, to quantify this contribution, a canonical analysis of principal coordinates was performed with the function *capscale* of the *vegan* package. Changes in relative abundance of ASV between the different seed samples were determined using linear discriminant analysis (LDA) effect size using the LefSe tool (Segata et al. 2011) available at http://huttenhower.sph.harvard.edu/galaxy.

To compare the beta dispersion amongst pollination modes, we quantified the variability in ASV composition within each pollination treatment using the betadisper function in the *vegan* package in R. Beta dispersion is measured by the distance to the centroid of each treatment group in the principal coordinate space (Anderson et al. 2006).

### Data availability

The raw sequencing data is available at the European Nucleotide Archive (ENA) under the study accession PRJEB31847. Tables and scripts used in this work are publicly available in GitHub.

## Acknowledgements

We thank Cédric Alaux, Stan Chabert, Cindy Morris, Ignasi Bartomeus for their input in the experimental design and interpretation of the results. Marianne Cousin for her help in the lab and Martial Briand for his help with DADA2 and other bioinformatic analyses. We also want to apologize to any authors whose relevant work was not cited in this article due to space constraints.

## Authors’ contributions

AP, BV, MB and GTC designed the study and made substantial contributions to the analysis and interpretation of the results. AP and GTC carried out the data analysis. AP and BV performed the tunnel experiments. GTC and BM constructed the amplicon libraries. AP and GTC wrote the manuscript with input from the other authors. All authors read and approved the final manuscript.

